# Transient 5-azacytidine exposure elicits persistent myogenic-like transcriptional remodeling in zebrafish fibroblasts: implications for cultivated seafood

**DOI:** 10.64898/2026.10.01.756133

**Authors:** Daisuke Ikeda, Nanami Mizusawa, Shugo Watabe

**Author notes:** Department of Biosphere Life Sciences, Graduate School of Bioresources, Mie University, 1577 Kurimamachiya-cho, Tsu, Mie 514-8507, Japan (Nanami Mizusawa). Corresponding author. E-mail address (D. Ikeda).

## Abstract

Fibroblasts are readily obtained and expanded and may provide an alternative cell source for cultivated seafood if their latent myogenic potential can be accessed. We examined whether transient exposure to 5-azacytidine (AZA), a cytidine analogue known to convert mouse embryonic fibroblasts into myoblasts, elicits a persistent muscle gene program in embryo-derived zebrafish (*Danio rerio*) Z3 fibroblasts. Cells were exposed to 0, 2.5, 5 or 10 µM AZA for 48 h and maintained in drug-free medium for 3 weeks. RNA-seq identified 744 genes that increased and 670 that decreased consistently across the concentration series. Increased genes encompassed myogenic regulators, fusion factors, sarcomeric components and excitation–contraction machinery, whereas decreased genes were enriched for extracellular-matrix and adhesion functions. In a separate six-dose RT-qPCR experiment (0–20 µM AZA) with three biological replicates per concentration, all 19 selected muscle genes were significantly increased at 1.25 µM, the lowest concentration tested, whereas 9 of 12 extracellular-matrix/fibroblast genes were significantly decreased. Several muscle genes plateaued or declined at higher concentrations, and multinucleated myotubes were not observed. A separate 5-aza-2′-deoxycytidine (decitabine) series showed a comparable muscle-associated response at 0.25 µM. These findings indicate that zebrafish fibroblasts retain access to a myogenic-like transcriptional program, and they provide a model for identifying food-compatible approaches to control fish-cell lineage plasticity for cultivated seafood.

## 1. Introduction

The ability of cytidine analogues to alter fibroblast identity is a foundational observation in skeletal-muscle biology. Exposure of non-myogenic mouse fibroblasts to 5-azacytidine (AZA) generated contractile striated-muscle cells as well as adipogenic and chondrogenic phenotypes [1,2]. These phenotypes appeared only several days or weeks after treatment, implying that cell division was required for their expression [2]. Subsequent analogue-comparison studies linked phenotype conversion to inhibition of DNA methylation [3], and myoblasts derived from AZA-treated C3H10T1/2 fibroblasts were later used to isolate MyoD, a single factor capable of converting fibroblasts into stable myoblasts [4].

DNA methylation participates in the regulation of myogenic genes. In mouse embryos, the *myogenin* promoter is methylated before activation and subsequently becomes demethylated [5], and muscle-specific demethylation accompanies lineage specification [6]. In Senegalese sole (*Solea senegalensis*) larvae, rearing temperature affects methylation of the putative *myogenin* promoter together with *myogenin* expression and muscle cellularity [7]. AZA is also active in fish cells: exposure of a zebrafish liver cell line to 0.5 µM AZA for 96 h caused promoter hypomethylation [8]. 5-Aza-2′-deoxycytidine (decitabine), a deoxycytidine analogue that is incorporated into DNA more directly than AZA, is also widely used to inhibit DNA methylation [9,10].

Whether comparable plasticity can be accessed in established fish fibroblast cultures remains poorly understood. Fish cell lines are valuable experimental resources, and robust, lineage-defined cell sources remain a practical constraint for cultivated seafood [11,12]. In our previous work, fibroblast-like cells progressively replaced myogenic cells during passage of Japanese eel (*Anguilla japonica*) muscle cultures, necessitating single-cell cloning to establish a stable myoblast line [13]. We subsequently established eel preadipocyte lines, further illustrating the need for complementary muscle, fat and connective-tissue cell resources [14].

Fin-derived fish cells have recently been explored as alternative starting material for cultivated seafood. Du et al. established a caudal-fin cell line from topmouth culter (*Culter alburnus*) and found that AZA alone (5 µM for 3 days) increased *myf5* expression but did not significantly increase myosin heavy chain expression or produce desmin-positive multinucleated myotubes [15]. An optimized five-compound regimen containing AZA, LY411575, RepSox, CHIR99021 and forskolin, followed by low-serum culture, produced multinucleated myotubes. Whether AZA alone can establish a broader and persistent myogenic program in fish fibroblasts after drug withdrawal therefore remained unclear.

Zebrafish is not a target species for cultivated seafood, but it is a widely used teleost model with a high-quality, well-annotated reference genome [16]. We therefore used zebrafish cells to establish a basis for future work in food fish species. Here, we asked whether AZA alone could elicit a persistent muscle gene program in established zebrafish Z3 fibroblasts. We examined the transcriptome 3 weeks after a 48-h AZA exposure by RNA-seq and functional enrichment analysis and confirmed selected responses in a separate six-dose RT-qPCR experiment. We also examined whether decitabine elicited comparable responses.

## 2. Materials and methods

### 2.1. Z3 cell culture and treatment with AZA and decitabine

Z3 cells, an embryo-derived zebrafish fibroblast culture established from the wild-type TL strain, were kindly provided by Prof Jun Hirayama (Bunkyo University, Japan). Related zebrafish embryo-derived cultures and their maintenance have been described previously [17]. The sex of the embryos from which the cells were derived was not determined. Cells were maintained at 28 °C in Leibovitz’s L-15 medium (Wako Pure Chemical Industries, Ltd., Osaka, Japan) supplemented with 10% fetal bovine serum and 1% antibiotic–antimycotic solution (Nacalai Tesque, Kyoto, Japan). AZA was obtained from Wako Pure Chemical Industries and dissolved in water. Cells were exposed to 0, 2.5, 5 or 10 µM AZA for 48 h. The medium was then replaced with drug-free growth medium, and cells were maintained for a further 3 weeks without passaging, with medium replacement every 3 days before RNA isolation. This post-exposure interval was selected because differentiated phenotypes in classical AZA-treated fibroblast cultures emerged over several days to weeks after transient exposure [2,3]. In a separate experiment, cells were exposed to 0, 0.25, 0.5 or 1.0 µM 5-aza-2′-deoxycytidine (decitabine; Wako Pure Chemical Industries), dissolved in water, using the same 48-h exposure and 3-week drug-free culture schedule.

### 2.2. RNA preparation, pooling, library construction and sequencing

For each AZA and decitabine concentration, total RNA was extracted separately from four independently cultured samples using ISOSPIN Cell & Tissue RNA (Nippon Gene, Tokyo, Japan) according to the manufacturer’s instructions. Equal amounts of RNA from the four samples were combined before library preparation to generate one composite RNA-seq library per concentration. Stranded cDNA libraries were prepared using the KAPA Stranded RNA-Seq Library Preparation Kit (Roche, Basel, Switzerland) according to the manufacturer’s instructions and sequenced in-house on an Illumina MiSeq using a MiSeq Reagent Kit v3 (Illumina, Inc., San Diego, CA, USA) in a 2 × 75-bp paired-end configuration.

### 2.3. RNA-seq processing and expression analysis

Raw paired-end reads were quality checked with FastQC, aligned to the zebrafish GRCz11 assembly with HISAT2 using Ensembl release 116 splice annotations, and summarized as reverse-stranded paired-end gene counts with featureCounts [18,19]. Detailed software versions and counting parameters are provided in Supplementary Methods.

Within the four-library Z3 AZA series, counts were normalized with DESeq2, and variance-stabilized values were used for sample correlation [20]. Because one composite library represented each concentration, expression responses were defined descriptively by effect size and directional consistency rather than by replicate-based significance testing. The principal response lists required an endpoint absolute log2 fold change of at least 1 and the same direction of change at all treated concentrations. An absolute log2 fold change of at least 2 was used as a sensitivity analysis. The decitabine series was normalized and analyzed separately against its own untreated library. Exact filters and transformations are reported in Supplementary Methods.

### 2.4. Functional enrichment analyses

Functional interpretation combined Gene Ontology over-representation analysis against the expressed-gene background, preranked gene-set enrichment analysis using complete gene rankings [21,22], and Metascape analysis [23]. The same functional analyses were applied to the decitabine series.

### 2.5. RT-qPCR and statistical analysis

Total RNA was extracted using ISOGEN II (Nippon Gene) and reverse transcribed with ReverTra Ace qPCR RT Master Mix (TOYOBO, Osaka, Japan). Real-time PCR was performed with THUNDERBIRD SYBR qPCR Mix (TOYOBO) on a LightCycler 96 instrument (Roche), following the manufacturer’s instructions and our published workflow [24]. In a separate experiment using the same 48-h exposure and 3-week drug-free culture schedule, cells were exposed to 0, 1.25, 2.5, 5, 10 or 20 µM AZA, with three independently seeded and treated cultures per concentration; each value represents the mean of technical duplicate reactions. Myogenic and extracellular-matrix/fibroblast genes were selected from the RNA-seq results and biological annotations (primer sequences in Supplementary Table S1).

Expression was normalized to elongation factor 1α (EF1α) and analyzed as ΔCq (Cq_target_ − Cq_EF1α_) by one-way analysis of variance followed by two-sided Dunnett comparisons of each AZA concentration with 0 µM. Relative expression (2^−ΔΔCq^), normalized to the untreated-group mean, is shown for visualization.

## 3. Results

### 3.1. AZA induced concentration-dependent transcriptome changes in Z3 cells

To examine the transcriptional state after transient AZA exposure, we sequenced Z3 cells 3 weeks after a 48-h treatment with 0, 2.5, 5 or 10 µM AZA. The four libraries contained 2.7–3.6 million read pairs, of which 91.7–91.9% aligned to the reference genome (Supplementary Table S2). Correlation with the untreated library decreased across the 2.5, 5 and 10 µM libraries (Pearson’s r = 0.96, 0.95 and 0.93, respectively), whereas the 5 and 10 µM libraries were the most similar pair (r = 0.98; Supplementary Fig. S1). Using a twofold threshold at 10 µM and a consistent direction of change at all treated concentrations, we identified 744 increased and 670 decreased genes. A stricter fourfold threshold retained 300 increased and 122 decreased genes with the same functional themes.

Three weeks after exposure, cultures treated with 5 or 10 µM AZA contained focal raised or ridge-like cell aggregates, whereas the untreated and 2.5 µM cultures showed no conspicuous morphological change (Fig. 1). No unequivocal multinucleated myotubes were observed.

**Figure 1.**
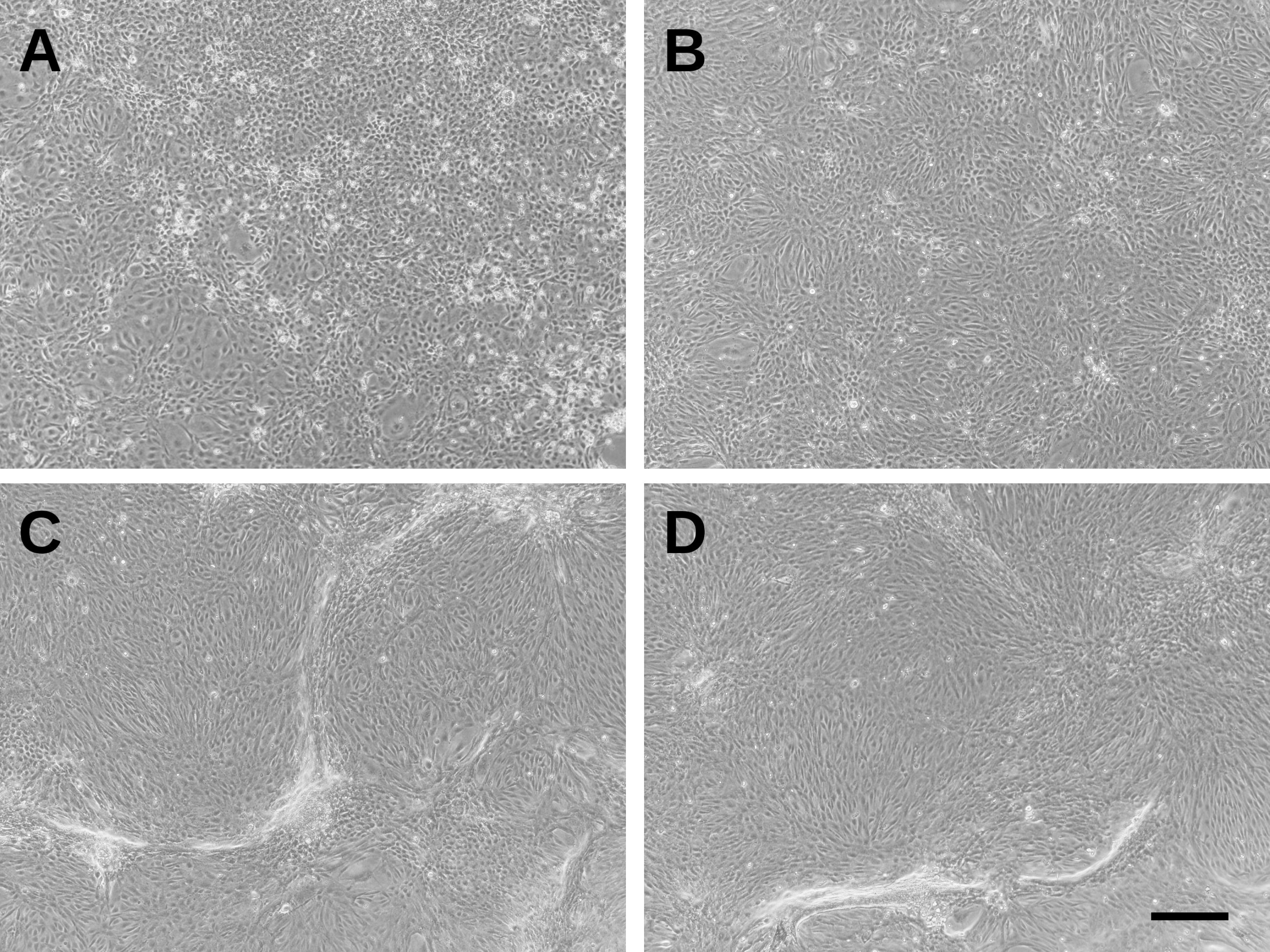
Morphology of Z3 cells 3 weeks after transient 5-azacytidine exposure. Representative phase-contrast micrographs of Z3 cells cultured with 0 (A), 2.5 (B), 5 (C), or 10 µM (D) 5-azacytidine (AZA) for 48 h. After AZA removal, the cells were maintained in growth medium for 3 weeks. Cells exposed to 0 or 2.5 µM AZA showed no marked morphological differences, whereas focal areas of raised, ridge-like cellular aggregation were observed in cultures treated with 5 or 10 µM AZA. However, no unequivocal morphological evidence of myogenic differentiation, such as multinucleated myotube formation, was observed. Scale bar, 200 µm.

### 3.2. AZA increased muscle gene expression and reduced extracellular-matrix gene expression

The increased genes included myogenic regulators and effectors spanning lineage specification (*myf5*, *myod1* and *myog*), cell fusion (*mymk* and *mymx*), sarcomeric structure and contraction (including *tnnt3a*, *ttn.1*, *ttn.2*, *desma* and *myhz2*), membrane/cytoskeletal organization (*dmd* and *klhl41a*), and excitation–contraction coupling (*atp2a1*, *tmem38a* and *ryr1b*). In contrast, the decreased genes included extracellular-matrix (ECM), adhesion and fibroblast-associated genes such as *ccn2b*, *col4a5*, *col2a1b*, *fn1b*, *lum*, *loxl2b*, *gpc4*, *adamtsl3*, *slit2* and *wif1*.

Untreated Z3 cells expressed ECM genes including *col1a1a*, *col1a1b*, *lum* and *fn1b*, while basal expression of the myogenic regulators and fusion genes *myf5*, *myod1*, *myog*, *mymk* and *mymx* was also detectable (Supplementary Table S7). The muscle-progenitor markers *pax7a* and *pax7b* did not pass the expression filter and showed only 0–5 raw counts across the libraries (Supplementary Table S7). After AZA exposure, myogenic regulatory, fusion and structural genes increased together, whereas ECM genes declined, indicating a shift from a fibroblast-like toward a myogenic-like expression profile.

Gene Ontology analysis linked the increased genes to muscle-related biological processes, including muscle system process (43 genes; 8.4-fold enrichment; false-discovery rate [FDR] = 2.5 × 10^−28^) and myofibril assembly (38 genes; 6.9-fold enrichment; FDR = 6.0 × 10^−21^). The decreased genes were associated with extracellular-matrix organization (25 genes; 4.3-fold enrichment; FDR = 1.9 × 10^−7^) and cell adhesion (53 genes; 3.3-fold enrichment; FDR = 6.9 × 10^−12^). Representative non-redundant biological-process terms are shown in Fig. 2, and complete results, including cellular-component and molecular-function ontologies, are provided in Supplementary Table S3.

**Figure 2.**
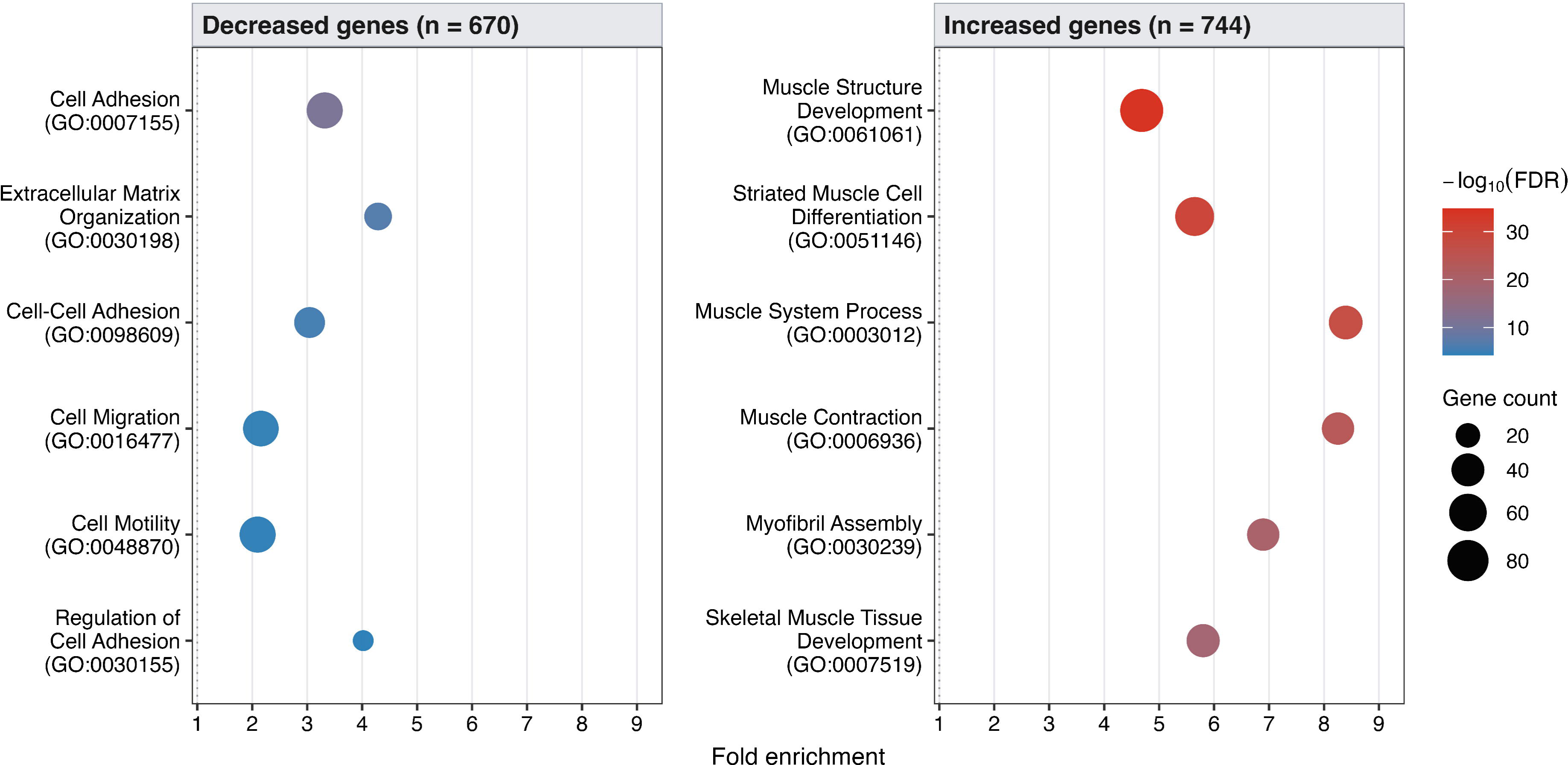
Gene Ontology biological-process over-representation analysis of AZA-responsive genes in Z3 cells. Terms are shown separately for increased and decreased genes. The expressed-gene background was used, and multiplicity was controlled by the Benjamini–Hochberg procedure. Redundant terms were reduced for presentation; the complete term-level results are provided in Supplementary Table S3.

Preranked gene-set enrichment analysis, which does not depend on a fold-change cutoff, gave the same result. Contractile-muscle-fiber gene sets were positively enriched at 2.5, 5 and 10 µM AZA (normalized enrichment score [NES] = 3.02, 2.95 and 2.98, respectively), whereas extracellular-matrix organization was negatively enriched (NES = −2.04, −2.29 and −2.49, respectively). Thus, the muscle response was already evident at 2.5 µM and changed little between 5 and 10 µM, whereas the ECM decrease became stronger with increasing concentration.

Metascape analysis yielded the same themes: muscle structure, differentiation, contraction and myofibril assembly for the increased genes, and extracellular-structure organization and cell adhesion for the decreased genes. These themes were also recovered from the more stringent sensitivity lists (Supplementary Table S3).

### 3.3. RT-qPCR confirmed the response in independent cultures from 1.25 µM AZA

To confirm selected RNA-seq results in independent cultures and to examine a wider concentration range, we performed RT-qPCR in a separate six-dose experiment (0–20 µM; three biological replicates per concentration). At 1.25 µM AZA, all 19 selected muscle genes were significantly increased, whereas 9 of 12 selected ECM/fibroblast genes were significantly decreased in Dunnett comparisons with the untreated group. Representative increases included *myf5* (2.1-fold), *mymk* (5.5-fold), *mymx* (4.0-fold), *myog* (7.5-fold) and *myhz2* (8.8-fold; Fig. 3; Supplementary Fig. S2 and Supplementary Table S4).

**Figure 3.**
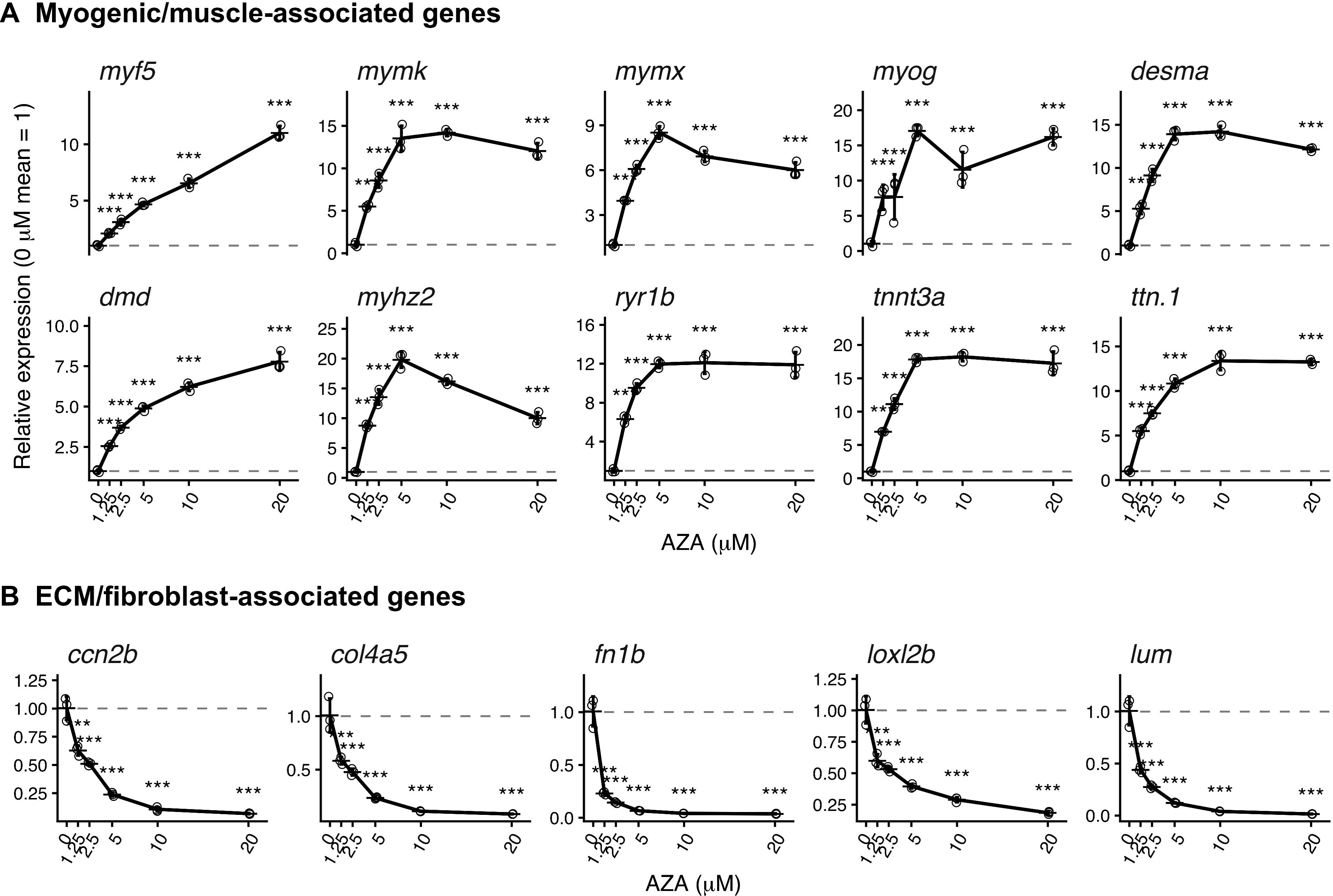
Six-dose RT-qPCR profiles of 15 representative genes in Z3 cells after transient AZA exposure. Panel A shows myogenic and muscle-associated genes; panel B shows extracellular-matrix/fibroblast-associated genes. Open circles represent independent biological replicates (n = 3 per concentration). Short horizontal strokes and error bars denote mean ± SD of 2^−ΔΔCq^ values normalized to the untreated-group mean; solid lines connect the group means. Statistical testing was performed on ΔCq values using one-way ANOVA followed by two-sided Dunnett comparisons with 0 µM. \**P* < 0.05, \*\**P* < 0.01, \*\*\**P* < 0.001. The remaining 16 genes are shown in Supplementary Fig. S2.

The muscle response was therefore detectable at 1.25 µM, which is lower than the lowest concentration examined by RNA-seq.. Several targets plateaued or declined at 20 µM; thus, the highest concentration did not necessarily produce the strongest response. *myod1* and *myog* also showed non-linear profiles, with prominent responses at intermediate concentrations. The four concentrations shared between RNA-seq and RT-qPCR showed broad descriptive agreement in response direction and profile shape (Supplementary Fig. S3).

### 3.4. Decitabine induced muscle genes at low concentration and a distinct series-wide response

In a separate experiment, Z3 cells were exposed to 0, 0.25, 0.5 or 1.0 µM decitabine using the same 48-h exposure and 3-week drug-free culture schedule as in the AZA experiment. At 0.25 µM, 343 genes met the criteria for low-dose peak-up genes and 353 for low-dose trough-down genes. The peak-up set was enriched for cytoskeleton in muscle cells (47 genes; 9.7-fold enrichment; q ≈ 1 × 10^−29^), muscle structure development (42 genes; 5.9-fold enrichment; q ≈ 1 × 10^−17^) and myofibril assembly (17 genes; 10.0-fold enrichment; q ≈ 2.5 × 10^−10^). A higher-confidence subset of 89 genes retained enrichment for muscle-related terms. Across the series, 358 genes that increased consistently were enriched for cell-cycle processes (42 of 314 annotated genes; 3.8-fold enrichment; q ≈ 1.6 × 10^−10^) and 467 genes that decreased consistently were enriched for extracellular-matrix organization (24 of 379 annotated genes; 6.7-fold enrichment; q ≈ 1.6 × 10^−10^; Supplementary Fig. S4 and Supplementary Tables S5 and S6).

## 4. Discussion

Transient AZA exposure was followed by persistent myogenic-like transcriptional remodeling in zebrafish Z3 fibroblasts. Here, “persistent” refers to a response that remained detectable 3 weeks after drug withdrawal, the only time point examined. The response extended from myogenic regulators and fusion genes to sarcomeric, membrane, excitation–contraction coupling and contraction machinery, while a fibroblast-associated extracellular-matrix and adhesion program declined. The present study characterized this response genome-wide after an extended drug-free interval, with AZA as the sole inducing compound.

The findings are consistent with the historical progression from AZA-induced phenotype switching to identification of MyoD [1–4] and suggest that an established teleost fibroblast culture retains access to a myogenic-like transcriptional program. Du et al. found that AZA alone increased *myf5* but did not significantly increase myosin heavy chain expression in forskolin-treated fin-derived cells after a 3-day exposure, whereas an AZA-containing five-compound regimen followed by low-serum culture produced desmin-positive multinucleated myotubes [13]. In Z3 cells examined 3 weeks after drug withdrawal, AZA alone was associated with a broader program that included fusion factors and sarcomeric myosin transcripts. Differences in cell source and concentration may contribute to this contrast, but the elapsed time after exposure may be particularly relevant, because in classical fibroblast studies new phenotypes appeared only several days to weeks after transient AZA exposure [2,3]. Neither Du et al. nor the present study observed myotube formation with AZA alone; in the present study, cells were not transferred to differentiation-permissive conditions, so their capacity for terminal differentiation remains untested.

In the RT-qPCR series, muscle genes responded at 1.25 µM, which is lower than the lowest concentration used for RNA-seq, whereas several responses plateaued or declined at 20 µM. Because lower concentrations and functional differentiation were not examined, these data do not define a minimum effective or optimal concentration.

Mechanistically, AZA and decitabine have overlapping but distinct intracellular fates. AZA is incorporated predominantly into RNA, although a metabolized fraction can enter DNA, whereas decitabine is incorporated more directly into DNA [9,10]. DNA-incorporated azacytosine covalently traps DNA methyltransferases [9,10], whereas incorporation of AZA into RNA can reduce protein synthesis [25]. The decitabine concentration range used here was one order of magnitude lower than the AZA range, consistent with the finding that decitabine produces DNA-mediated effects at 2- to 10-fold lower concentrations than AZA in leukemia cell lines [25]. Notably, 0.25 µM decitabine produced a muscle-associated response with strong enrichment for muscle structure development and myofibril assembly, even though decitabine differs from AZA in its intracellular fate. That two cytidine analogues with different metabolism elicited similar muscle-related responses strengthens the observation, although their series-wide profiles differed. These data do not establish DNA demethylation as the cause and do not permit a direct comparison of potency or cytotoxicity. Although DNA methylation participates in myogenic gene regulation in mammals and fish [5–7] and AZA can cause promoter hypomethylation in zebrafish cells [8], these observations do not demonstrate the mechanism operating in Z3 cells. Decitabine can also cause growth arrest and a senescence-associated secretory phenotype in a fish cell line [26], and cell viability and stress responses were not measured here. The 3-week drug-free interval makes continued direct incorporation an unlikely explanation for the final profile, but the present study cannot distinguish a persistent regulatory change from selection or expansion of a responsive subpopulation.

The work has longer-term relevance to cellular agriculture and to the broader phenotypic plasticity of fish cell lines [27]. Fibroblast-like cells can overgrow primary fish muscle cultures and complicate preservation of myoblast identity, as observed in our Japanese eel studies [13,14]. Although AZA itself is unsuitable for food production, the present findings provide a basis for screening food-compatible compounds and culture conditions that may reproduce aspects of the myogenic-like response observed here. Vitamin C is one candidate, because it enhances myogenic differentiation of mouse C2C12 myoblasts through TET- and KDM7A-dependent demethylation pathways [28]. Future studies should determine whether vitamin C or other food-compatible modulators, alone or combined with differentiation-promoting culture conditions, can induce a comparable response and support terminal differentiation in fish fibroblasts. The Z3 system therefore serves as a model for identifying regulatory pathways rather than as a direct protocol for producing edible tissue.

Several limitations should be considered. First, each RNA-seq concentration was represented by one composite library generated by pooling RNA from four independent cultures. Pooling reduced the influence of any single culture but did not preserve information on between-culture variability. The RNA-seq analysis therefore describes effect sizes, concentration-associated patterns and functional enrichment rather than statistically significant gene-level differential expression. The separate six-dose RT-qPCR experiment provided biological-replicate support for selected genes but did not provide transcriptome-wide replication. Second, EF1α was used as the sole RT-qPCR reference gene. Its mean Cq varied by no more than 0.48 cycles across the six concentrations, providing descriptive evidence of limited variation in this dataset but not formal reference-gene validation. Third, protein abundance, DNA methylation, chromatin accessibility, cell viability, proliferation and functional contractile differentiation were not measured, and cells were maintained in growth medium containing 10% fetal bovine serum rather than differentiation medium. Finally, the design cannot distinguish reprogramming of individual fibroblasts from selection or expansion of a pre-existing responsive subpopulation.

In conclusion, transient AZA exposure was followed by persistent, concentration-dependent myogenic-like transcriptional remodeling in zebrafish embryo-derived fibroblasts, comprising induction of myogenic regulators, fusion factors and sarcomeric genes and attenuation of extracellular-matrix programs, with muscle gene increases detectable from 1.25 µM AZA. A comparable response was observed with decitabine at low concentration. These findings indicate that an established fish fibroblast culture retains access to a myogenic-like transcriptional program and provide a model for identifying food-compatible approaches to controlling fish-cell lineage plasticity for cultivated seafood.

## CRediT authorship contribution statement

Daisuke Ikeda: Conceptualization, Data curation, Formal analysis, Funding acquisition, Investigation, Methodology, Project administration, Supervision, Visualization, Writing – original draft, Writing – review & editing. Nanami Mizusawa: Investigation, Writing – review & editing. Shugo Watabe: Conceptualization, Writing – review & editing.

## Data availability

Raw RNA-seq reads from the Z3 AZA and decitabine series have been deposited in the DDBJ Sequence Read Archive under BioProject accession number PRJDB45773.

## Declaration of generative AI and AI-assisted technologies in the manuscript preparation process

During the preparation of this work, the authors used OpenAI Codex to assist in drafting analysis scripts, and Anthropic Claude for language editing and manuscript review. After using these tools, the authors reviewed and edited the content as needed and take full responsibility for the content of the published article.

## Acknowledgments

We thank Prof Jun Hirayama (Bunkyo University, Japan) for providing the Z3 cells and Natsuyuki Kanatani and Yuto Ishikawa for assistance with the experiments.

