## Supplementary Methods for "Transient 5-azacytidine exposure elicits persistent myogenic-like transcriptional remodeling in zebrafish fibroblasts: implications for cultivated seafood"

**S1. Sequencing, reference files, read quality control, alignment and counting**

Libraries were sequenced using the MiSeq Reagent Kit v3 (150 cycles). Reads were analyzed against the Danio rerio GRCz11 primary assembly using the matched Ensembl release 116 annotation (Danio_rerio.GRCz11.116.gtf). Raw paired FASTQ files were evaluated with FastQC. No additional read trimming was performed. Reads were aligned with HISAT2 using splice sites extracted from the matched GTF. Coordinate-sorted and indexed BAM files were generated with samtools and checked with samtools quickcheck. Library orientation was evaluated with RSeQC. Gene-level fragments were counted with featureCounts against annotated exons grouped by gene_id, using paired-end, properly paired, non-chimeric and reverse-stranded counting.

**S2. Normalization and descriptive expression-change definitions**

The four Z3 AZA libraries were normalized as one series with DESeq2. A blind variance-stabilizing transformation was used for Pearson correlation and sample-level visualization. Genes with at least 10 raw fragments in at least two libraries were retained as expressed. Endpoint effect size was calculated from size-factor-normalized counts as log2[(normalized count at 10 µM + 1)/(normalized count at 0 µM + 1)]. Because each concentration was represented by one composite library, replicate-based gene-level P values, adjusted P values and false-discovery-rate thresholds were not used to define differentially expressed genes.

The principal increase and decrease lists required an absolute endpoint log2 fold change of at least 1 at 10 µM versus 0 µM and the same response direction at 2.5, 5 and 10 µM. These lists contained 744 increased and 670 decreased genes. A fourfold endpoint threshold (absolute log2 fold change of at least 2) yielded 300 increased and 122 decreased genes as a sensitivity analysis.

**S3. Functional enrichment analysis**

Gene Ontology (GO) over-representation analysis used the expressed-gene universe as the custom background. Increased and decreased response lists were analyzed separately for biological process, cellular component and molecular function. Hypergeometric P values were adjusted with the Benjamini–Hochberg procedure within each list and ontology. Representative non-redundant biological-process terms were selected for the main figure; complete term-level results are supplied in Supplementary Table S3.

Preranked gene-set enrichment analysis used the complete expressed-gene ranking for each treated concentration relative to the untreated library. Analyses were performed with clusterProfiler and the fgsea backend. Gene sets containing 15–500 genes were tested with exponent = 1 and eps = 0.

Metascape analyses were conducted with Danio rerio specified as both the input species and the analysis species and with the expressed-gene universe supplied as the background. Increased and decreased lists were analyzed separately and as multi-list submissions. Metascape q values refer to multiple-testing-adjusted enrichment probabilities.

**S4. Decitabine response analysis**

Genes with a raw fragment count of at least 10 in at least two of the four decitabine libraries were retained. Fold changes were calculated from DESeq2 size-factor-normalized counts after adding a pseudocount of 1. Low-dose peak-up genes had log2 fold change (0.25 µM versus 0 µM) ≥ 1 and a greater response at 0.25 µM than at both 0.5 and 1.0 µM; low-dose trough-down genes were defined analogously. Series-wide directionally consistent genes had an absolute endpoint log2 fold change (1.0 µM versus 0 µM) ≥ 1, with all three treated concentrations on the same side of the untreated value. A higher-confidence low-dose peak-up subset required log2 fold change ≥ 2 at 0.25 µM. The principal peak-up, trough-down, series-wide increase and series-wide decrease lists contained 343, 353, 358 and 467 genes, respectively; the higher-confidence peak-up subset contained 89 genes. These exploratory effect-size categories do not represent statistically significant differential expression because one composite library represented each concentration.

**S5. Software and resource summary**

Analysis scripts were drafted with the assistance of OpenAI Codex; all code was executed, inspected and verified by the authors using the software and resources documented in this section.

| Resource | Version/release | Purpose | Key setting |
| --- | --- | --- | --- |
| Danio rerio genome | GRCz11 | Read alignment | Primary assembly |
| Ensembl annotation | Release 116 | Splice sites and gene models | Matched to GRCz11 |
| FastQC | 0.12.1 | Raw-read quality control | Paired FASTQ files |
| HISAT2 | 2.2.1 | Splice-aware alignment | Matched Ensembl splice sites |
| samtools | 1.17 | BAM processing and integrity checks | Coordinate-sorted BAM |
| RSeQC | 5.0.4 | Library-orientation assessment | Strandedness check |
| featureCounts | 2.1.1 | Gene-level fragment counting | Paired-end; reverse-stranded |
| DESeq2 | 1.46.0 | Normalization and VST | No replicate-based DEG testing |
| clusterProfiler | 4.14.0 | GO analysis and GSEA | Expressed-gene background |
| fgsea | 1.32.2 | Preranked enrichment backend | eps = 0 |
| Metascape | Web analysis | Independent functional enrichment | D. rerio input and analysis species |
