## Supplementary Fig. for "Transient 5-azacytidine exposure elicits persistent myogenic-like transcriptional remodeling in zebrafish fibroblasts: implications for cultivated seafood"

### Slide 1
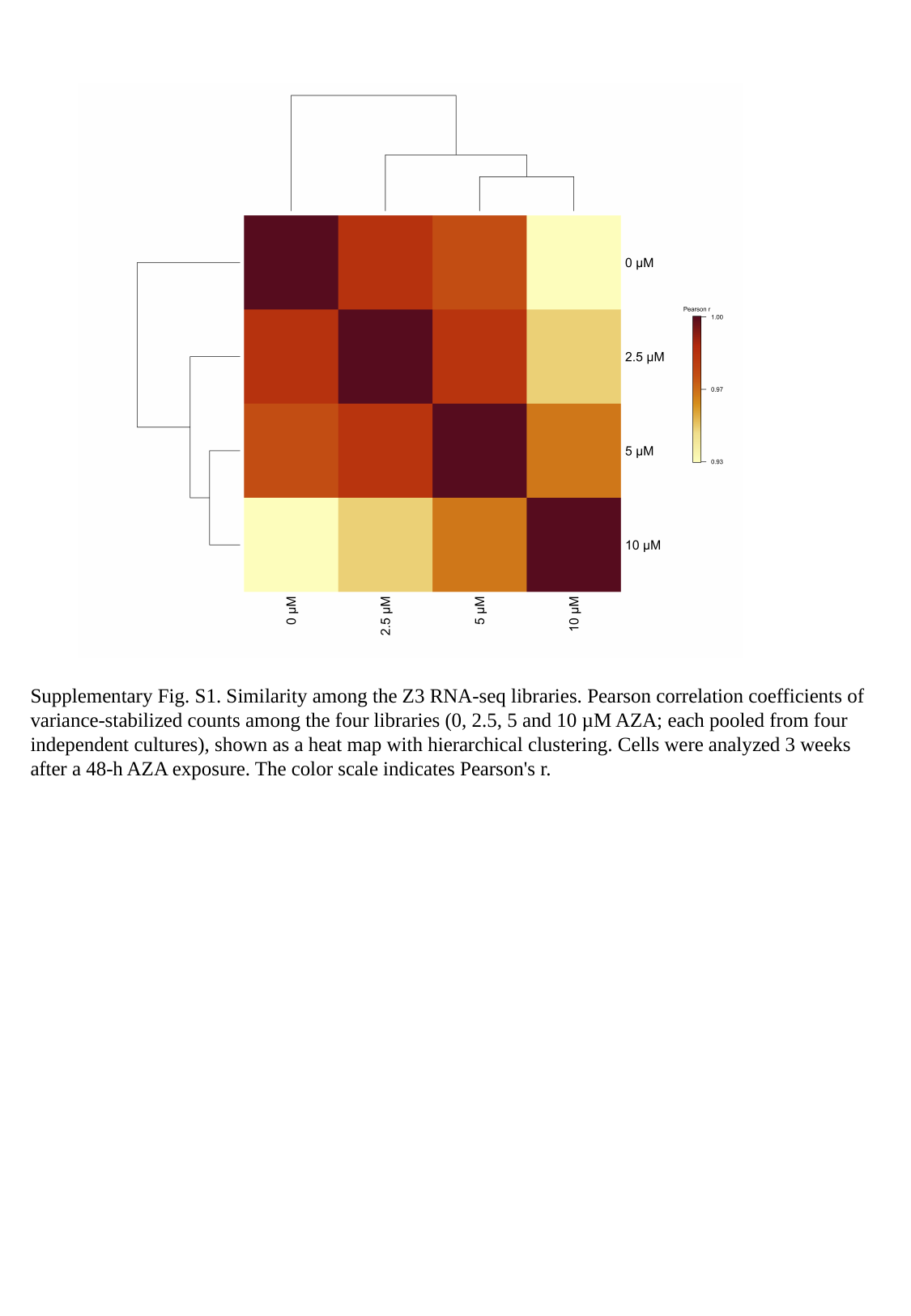

Supplementary Fig. S1. Similarity among the Z3 RNA-seq libraries. Pearson correlation coefficients of variance-stabilized counts among the four libraries (0, 2.5, 5 and 10 µM AZA; each pooled from four independent cultures), shown as a heat map with hierarchical clustering. Cells were analyzed 3 weeks after a 48-h AZA exposure. The color scale indicates Pearson's r.

### Slide 2
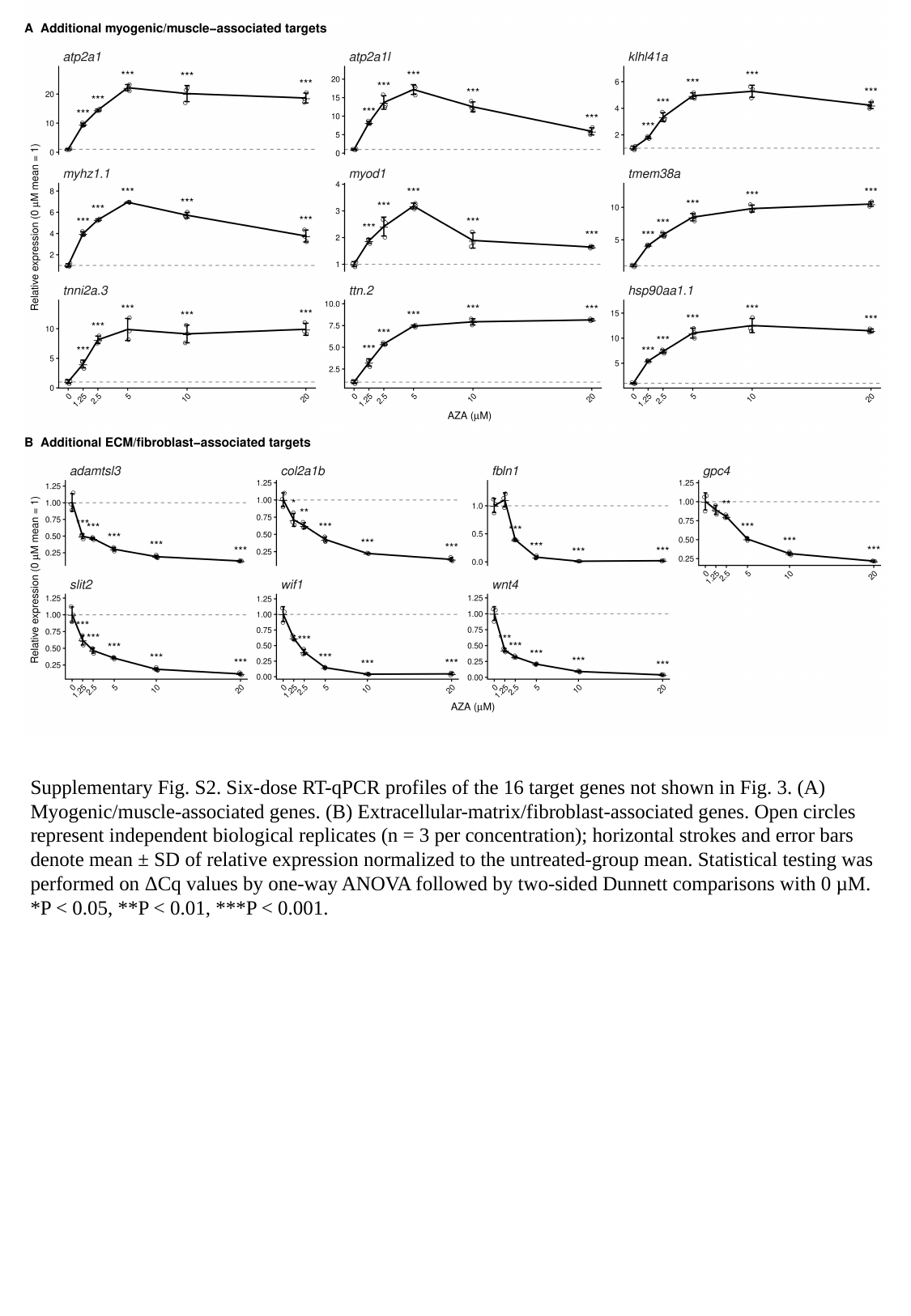

Supplementary Fig. S2. Six-dose RT-qPCR profiles of the 16 target genes not shown in Fig. 3. (A) Myogenic/muscle-associated genes. (B) Extracellular-matrix/fibroblast-associated genes. Open circles represent independent biological replicates (n = 3 per concentration); horizontal strokes and error bars denote mean ± SD of relative expression normalized to the untreated-group mean. Statistical testing was performed on ΔCq values by one-way ANOVA followed by two-sided Dunnett comparisons with 0 µM. *P < 0.05, **P < 0.01, ***P < 0.001.

### Slide 3
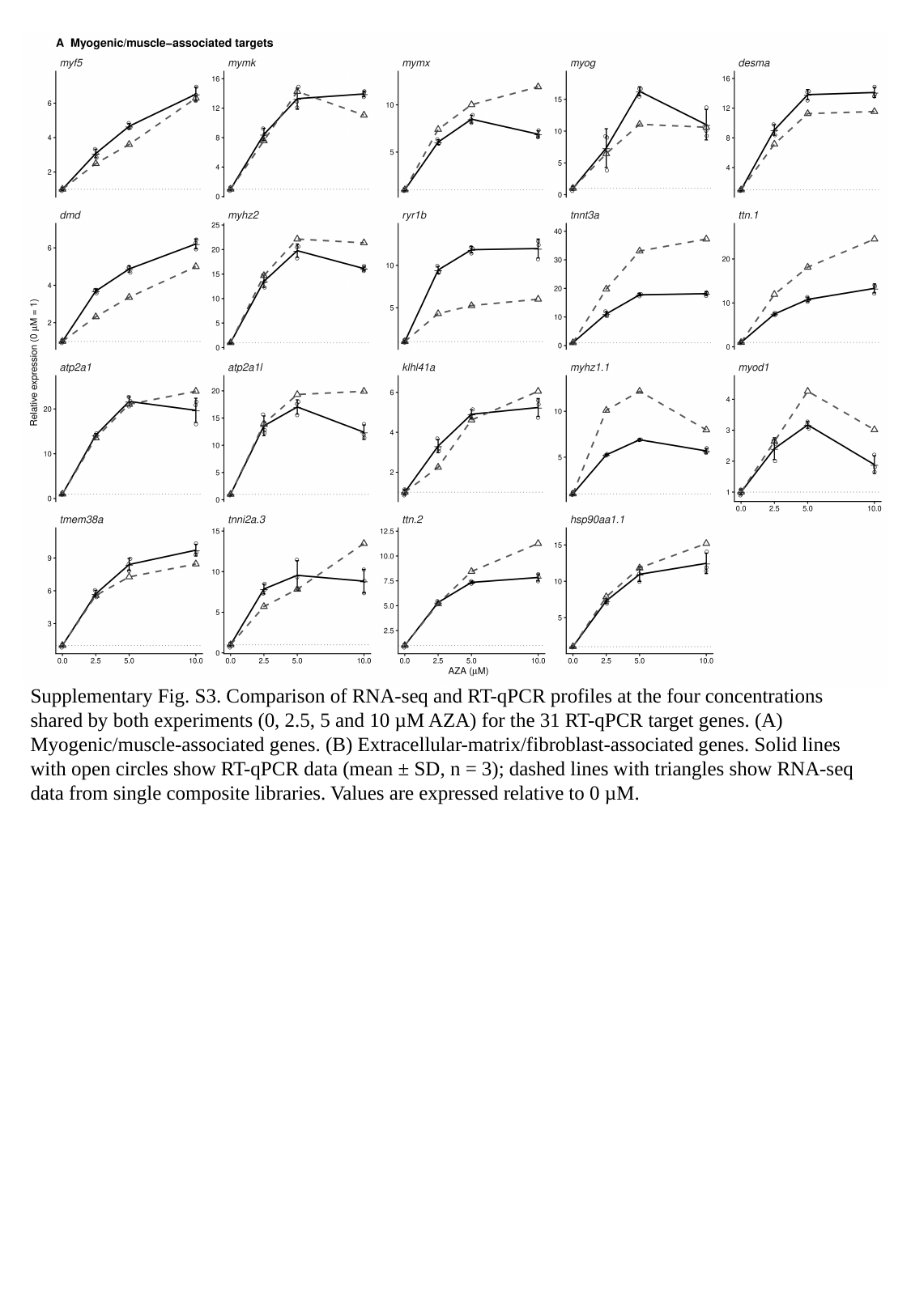

Supplementary Fig. S3. Comparison of RNA-seq and RT-qPCR profiles at the four concentrations shared by both experiments (0, 2.5, 5 and 10 µM AZA) for the 31 RT-qPCR target genes. (A) Myogenic/muscle-associated genes. (B) Extracellular-matrix/fibroblast-associated genes. Solid lines with open circles show RT-qPCR data (mean ± SD, n = 3); dashed lines with triangles show RNA-seq data from single composite libraries. Values are expressed relative to 0 µM.

### Slide 4
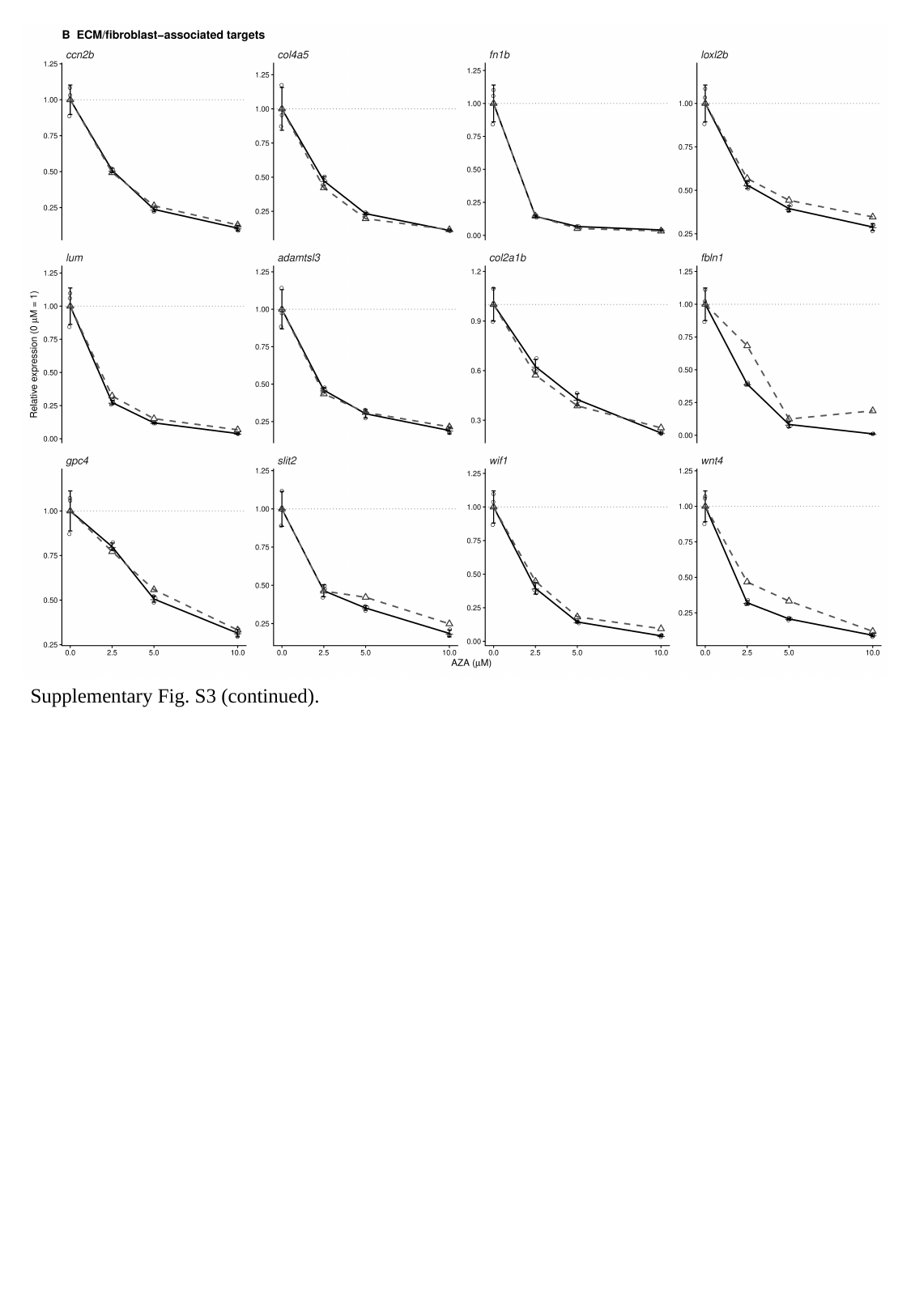

Supplementary Fig. S3 (continued).

### Slide 5
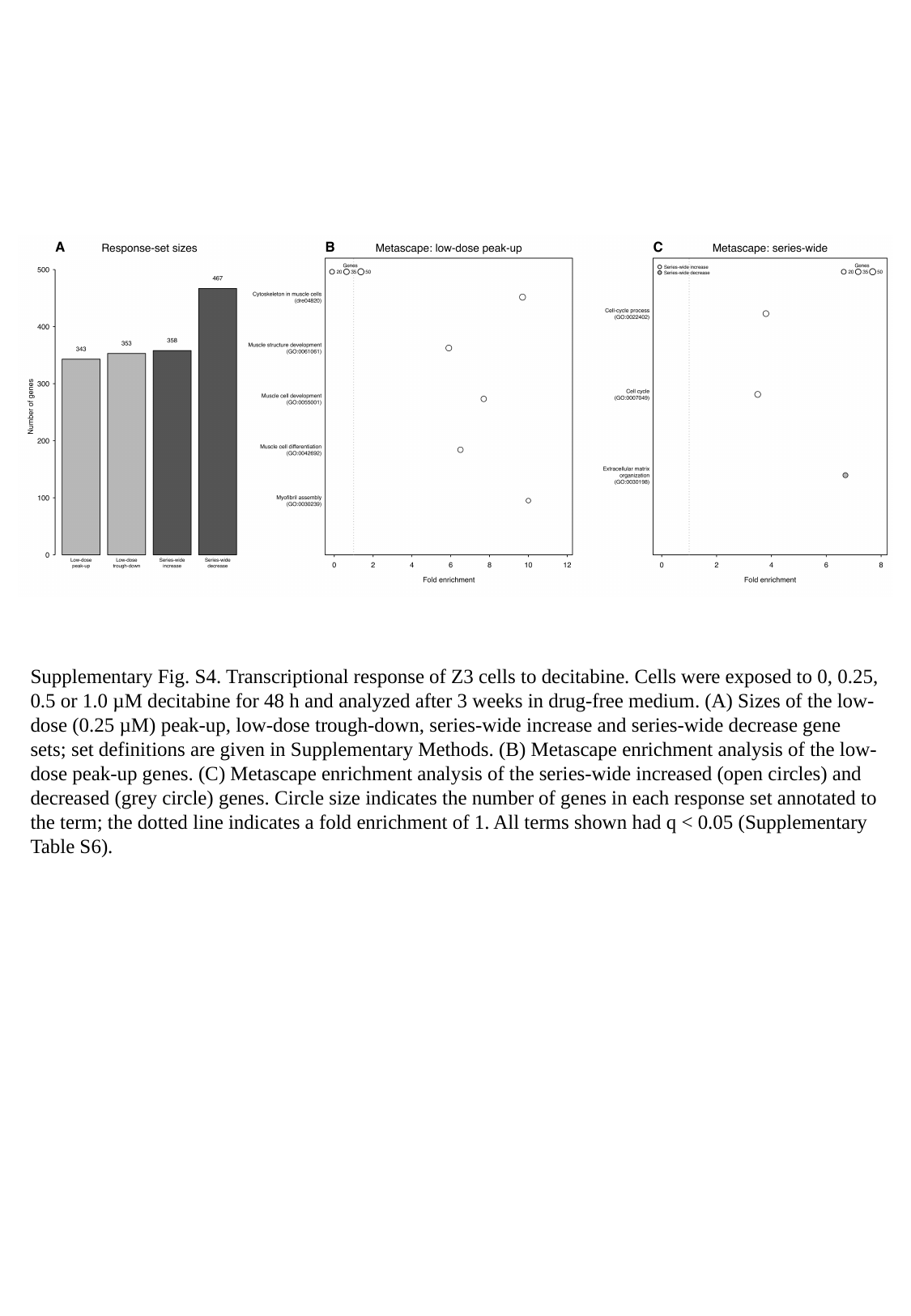

Supplementary Fig. S4. Transcriptional response of Z3 cells to decitabine. Cells were exposed to 0, 0.25, 0.5 or 1.0 µM decitabine for 48 h and analyzed after 3 weeks in drug-free medium. (A) Sizes of the low-dose (0.25 µM) peak-up, low-dose trough-down, series-wide increase and series-wide decrease gene sets; set definitions are given in Supplementary Methods. (B) Metascape enrichment analysis of the low-dose peak-up genes. (C) Metascape enrichment analysis of the series-wide increased (open circles) and decreased (grey circle) genes. Circle size indicates the number of genes in each response set annotated to the term; the dotted line indicates a fold enrichment of 1. All terms shown had q < 0.05 (Supplementary Table S6).
